# Asymmetric evolution of protein domains in the leucine-rich repeat receptor-like kinase (LRR-RLK) family of plant developmental coordinators

**DOI:** 10.1101/2023.03.13.532436

**Authors:** Jarrett Man, Thomas Harrington, Kyra Lally, Madelaine E. Bartlett

**Affiliations:** Department of Biology, University of Massachusetts Amherst; Amherst, MA 01002, USA

## Abstract

The coding sequences of developmental genes are expected to be conserved over deep time, with *cis-*regulatory change driving the modulation of gene function. In contrast, proteins with roles in defense are expected to evolve rapidly, in molecular arms-races with pathogens. However, some gene families include both developmental and defense genes. In these families, does the tempo and mode of evolution differ between developmental and defense genes, despite shared ancestry and structure? The leucine-rich repeat receptor-like kinase (LRR-RLKs) protein family includes many members with roles in plant development and defense, thus providing an ideal system for answering this question. LRR-RLKs are receptors that traverse plasma membranes. LRR domains bind extracellular ligands, RLK domains initiate intracellular signaling cascades in response to ligand binding. In LRR-RLKs with roles in defense, LRR domains evolve faster than RLK domains. To determine whether this asymmetry extends to developmental LRR-RLKs, we assessed evolutionary rates and tested for selection acting on eleven clades of LRR-RLK proteins, using deeply sampled protein trees. To assess functional evolution, we performed heterologous complementation assays using *Arabidopsis thaliana* (arabidopsis) LRR-RLK mutants. We found that the LRR domains of developmental LRR-RLK proteins evolved faster than their cognate RLK domains. LRR-RLKs with roles in development and defense had strikingly similar patterns of molecular evolution. Heterologous transformation experiments revealed that the evolution of developmental LRR-RLKs likely involves multiple mechanisms, including changes to *cis-*regulation, coding sequence evolution, and escape from adaptive conflict. Our results indicate similar evolutionary pressures acting on developmental and defense signaling proteins, despite divergent organismal functions. In addition, deep understanding of the molecular evolution of developmental receptors can help guide targeted genome engineering in agriculture.

## Introduction

Signaling from outside a cell to direct cellular behavior is critical in both response to pathogens and development. In defense signaling, proteins directly bind extracellular pathogen effectors to trigger intracellular defense processes. In development, similar extracellular signals bind receptors to trigger intracellular cell identity and physiological responses. These two functional categories of signaling proteins are often structurally similar, but for those with roles in defense, the effector-binding domains often evolve more rapidly and with greater rates of positive selection (X.S. Zhang et al. 2006; Fischer et al. 2016; Ahmad et al. 2021; Ghosh et al. 2022). Rapid evolution in defense ectodomains is a response to rapidly evolving pathogen effectors in a molecular arms-race, while intracellular domains still trigger conserved host reactions and may evolve slower (Lehti-Shiu et al. 2012). In contrast, for developmental receptors, it is a long standing conjecture that regulatory evolution is more important than changes to protein structure and function (King and Wilson 1975; Atchley and Hall 1991; Carroll 2008; Wittkopp and Kalay 2011; Long et al. 2016; Marand et al. 2023). The leucine-rich repeat receptor-like kinase (LRR-RLK) protein family of receptors has roles in both development and defense (Liu et al. 2017). These proteins have a conserved domain structure; the LRR domain is the ligand-binding ectodomain at their N terminus, the RLK domain is an intracellular kinase domain, with the two separated by a single-strand transmembrane α-helix (Shiu and Bleecker 2001). This domain structure within a protein family that orchestrates both defense and development allows direct comparisons of protein domain evolution. While *cis-*regulatory evolution is undoubtedly important, the evolution of protein structure and function may be an important driving force in the evolution of developmental signaling pathways (Hoekstra and Coyne 2007). Here, we investigate whether asymmetric evolution of extracellular signal perception and intracellular signal response domains is limited to defense proteins, or if the trend also applies to signaling proteins with roles in development.

There is suggestive evidence that sequence encoding LRR and RLK domains are evolving asymmetrically within the same developmental gene. First, although encoded by single genes, LRR and RLK domains are physically separated by the plasma membrane, and have distinct molecular functions (Santiago et al. 2016; Hohmann et al. 2018). Second, different clades of developmental LRR-RLKs respond to distinct ligands, but activate similar intracellular signaling cascades, implying asymmetric functional shifts (Hohmann et al. 2018; Zheng et al. 2019; Zhu et al. 2019; Liu et al. 2022). Third, the coordination between cognate LRR and RLK domains is interchangeable: functional output by RLK domains can be activated by alternate LRR domains in chimeric proteins (Osakabe et al. 2005; Diévart et al. 2006; Brutus et al. 2010; Zheng et al. 2019; Hohmann et al. 2020). This uncoupled domain function may also permit independent evolutionary trajectories (Bhattacharyya et al. 2006; Di Roberto and Peisajovich 2014; Sato et al. 2014). Fourth, over deep time, truncations or fusions with unrelated domains are more common in LRR domains than in RLK domains, and more LRR-only genes are conserved and expressed (Man et al. 2020). Taken together, these data suggest that developmental LRR and RLK domains may be evolving asymmetrically.

To determine whether asymmetric trends of evolution exist in the coding portions of developmental LRR-RLKs, we leveraged the One Thousand Plant Transcriptomes database (One Thousand Plant Transcriptomes Initiative, 2019) and inferred deeply sampled peptide trees for 11 subfamilies of LRR-RLK proteins (Shiu and Bleecker 2001; Dufayard et al. 2017; Man et al. 2020). We used these trees to test for distinct evolutionary forces acting on LRR vs. RLK domains. We also used heterologous transformation experiments to test for divergent functional evolution of LRR vs. RLK domains. We discovered a clear signal of asymmetric evolution between LRR and RLK domains encoded by the same gene, but our heterologous transformation experiments revealed more complexity. Our work highlights the multiple interacting mechanisms that drive the evolution of gene and protein function.

## Results

### LRR domains are evolving faster than RLK domains

To determine the evolutionary trajectories of individual protein domains, we needed deeply sampled trees that included multiple LRR-RLK clades with functions in development and defense. Therefore, we selected nine LRR-RLK sub-families in clade XI with roles in plant development and physiology, and two LRR-RLKs with roles in defense (Table 1). We also included CHITIN ELICITOR RECEPTOR KINASE 1 (CERK1), a defense receptor LysM-RLK that binds chitin derivatives (Chinchilla et al. 2006; Gimenez-Ibanez et al. 2009; Zhang et al. 2015) (Table 1). CERK1 binds pathogen-associated molecular patterns, is a target of bacterial effectors, and is expected to be under intense positive selection pressure (Gimenez-Ibanez et al. 2009; Hogenhout et al. 2009; Lohmann et al. 2010; De Mita et al. 2014; Wang et al. 2018).

**Table 1.** protein clades included in gene tree inference

| Name (*A. thaliana*) | Gene ID | Biological Function | Clade<sup>1</sup> | Genes<sup>2</sup> | Reference |
| --- | --- | --- | --- | --- | --- |
| *BARELY ANY MERISTEM (BAM1)* | AT5G65700 | Meristem specification | XI | 447 | (DeYoung et al. 2006) |
| *C-TERMINALLY ENCODED PEPTIDE RECEPTOR 2 (CEPR2)* | AT1G72180 | Nitrate uptake | XI | 358 | (Tabata et al. 2014) |
| *CHITIN ELICITOR RECEPTOR KINASE 1 (CERK1)* | AT3G21630 | Microbe detection | NA<sup>3</sup> | 367 | (Miya et al. 2007) |
| *CLAVATA1 (CLV1)* | AT1G75820 | Shoot meristem size | XI | 156 | (Clark et al. 1993) |
| *FLAGELLIN-SENSITIVE 2 (FLS2)* | AT5G46330 | Microbe detection | XII | 60 | (Gómez-Gómez and Boller 2000) |
| *GASSHO2 (GSO2)* | AT5G44700 | Epidermis development | XI | 17 | (Tsuwamoto et al. 2008) |
| *HAESA (HAE)* | AT4G28490 | Abscission, cell separation | XI | 409 | (Jinn et al. 2000) |
| *HAESA-LIKE3 (HSL3)* | AT5G25930 | Stomatal closure | XI | 208 | (Liu et al. 2020; Liu et al. 2022) |
| *HAIKU2 (IKU2)* | AT3G19700 | Seed, embryo development | XI | 354 | (Luo et al. 2005) |
| *PLANT ELICITOR PEPTIDE1 RECEPTOR1 (PEPR1)* | AT1G73080 | Innate immunity | XI | 41 | (Liu et al. 2013) |
| *PHLOEM INTERCALATED WITH XYLEM (PXY)* | AT5G61480 | Vascular development | XI | 137 | (Hirakawa et al. 2008) |
| *ROOT MERISTEM GROWTH FACTOR1 INSENSITIVE5 (RGI5)* | AT1G34110 | Root meristem size | XI | 8 | (Ou et al. 2016) |

<sup>1</sup>Clade assignments from (Man et al. 2020). <sup>2</sup>Number of genes recovered in each clade. <sup>3</sup>CERK1 is a LysM-RLK.

Gene trees with deep taxonomic sampling are needed to detect signatures of selection and variation in evolutionary rate. However, fully assembled plant genomes are relatively few given the immense diversity of multicellular land plants, and are not phylogenetically evenly distributed. This limits both bioinformatic resolution, and skews statistical inference towards overrepresented clades, like the grasses (Kress et al. 2022). To compensate for this sampling bias, we leveraged the One Thousand Plant Transcriptomes project (1KP), which includes phylogenetically informed sampling from across multicellular land plants (One Thousand Plant Transcriptomes Initiative 2019). Using the 1KP data and an iterative search and tree inference algorithm we developed (Man et al. 2020), we inferred deeply sampled peptide trees for all 11 LRR-RLK subfamilies, each of which had between 8 and 447 members (Table 1). This wide discrepancy in the number of genes recovered may be due to the 1KP sampling strategy, which focused preferentially on above-ground tissue, biasing against root-expressed genes like *RGI5 (Ou et al. 2016; One Thousand Plant Transcriptomes Initiative 2019)*.

To ask how the tempo of evolution might differ between LRR and RLK domains, we used our deeply sampled trees to estimate evolutionary rates of amino acid substitutions in angiosperm LRR-RLK sequences using multiple methods. First, we used our trees to infer site-specific relative substitution rates using an empirical Bayesian approach implemented in *IQ-TREE* (fig. 1) (Nguyen et al. 2015). This analysis permitted the comparison of amino acid site substitution rates in various regions of each protein, relative to overall mean substitution rates. For example, in the *CLAVATA1* (*CLV1*) clade, substitution rates were lowest in the region of the RLK domain (34.2% of the overall mean rate), substantially higher rates in the region of the LRR domain (95.7% of the overall mean rate), and the fastest rates were in inter-domain sequences. Despite the different functional roles of proteins in each clade (Table 1), these analyses revealed similar patterns of site-substitution rates between clades (fig. 1B and 1C). Averaged over all 11 independent clades, sequences of LRR domains evolve at 90.1% of the clade’s overall mean substitution rate, while sequences of RLK domains evolve slower at 66.7% of the mean (fig. 1E). We next employed a tangential approach; a Markov chain Monte Carlo Bayesian analysis implemented in *BEAST*, to infer the substitution rate for each domain that includes probability estimates of the entire tree structure (Bouckaert et al. 2014). This analysis revealed that LRR domains were evolving significantly faster than RLK domains in each clade (unpaired Wilcoxon tests, *p*-value < .001) (fig. 1D). Notably, while the relative rates of ectodomain evolution were substantially higher for CERK1 than for the nine developmental proteins we analyzed, this was not the case for FLS2 or PEPR1, which are middle of the pack with respect to asymmetric evolution rate. Therefore, the differences in evolutionary rates between LRR and RLK domains are not solely a function of role. Independent evolutionary rates for LRR and RLK domains may be consistent across LRR-RLKs.

**Figure 1:**
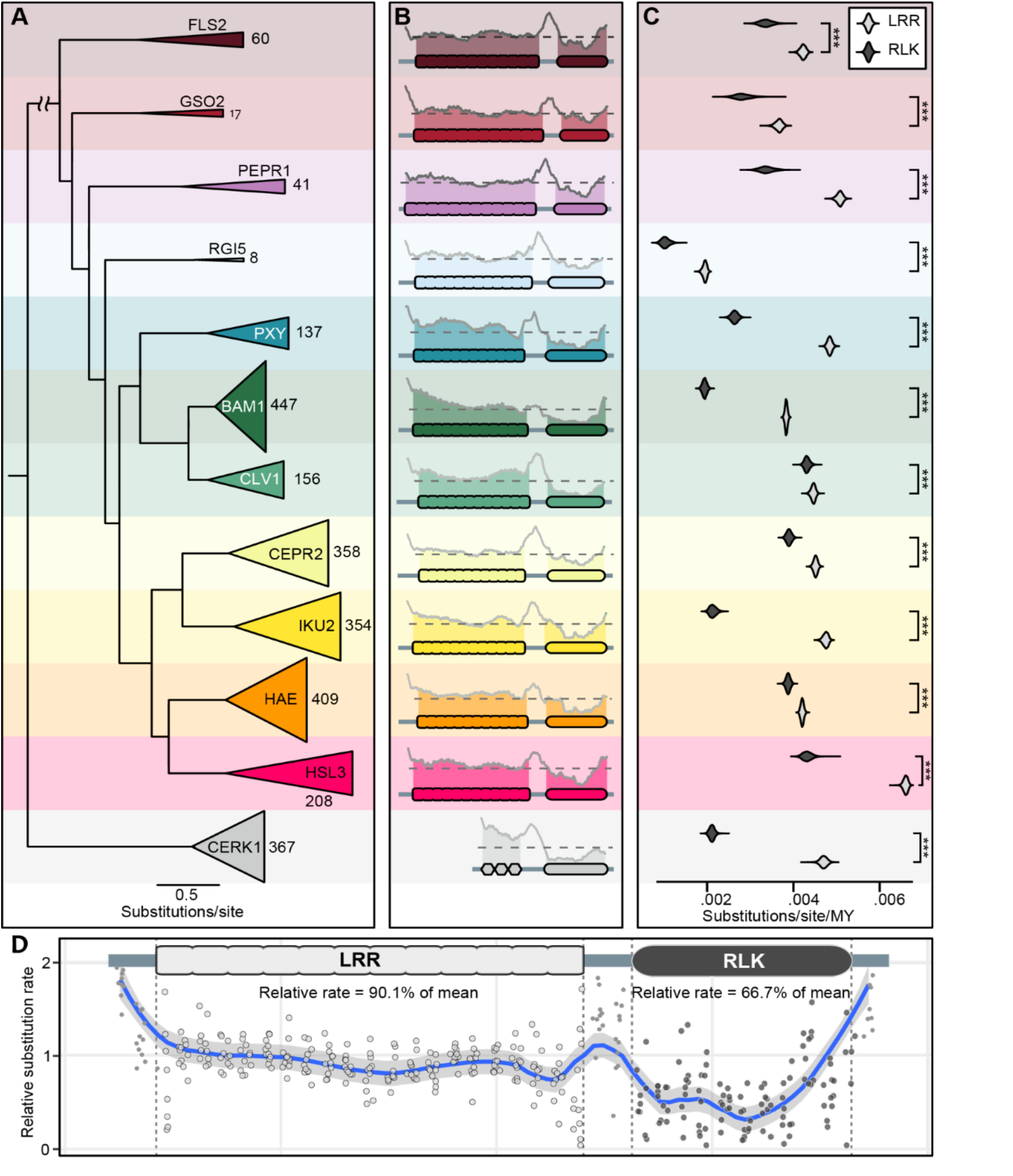
Evolutionary rates differ between LRR and RLK domains. *(A)* Relationships of 11 LRR-RLK clades and *CERK1*. Clade triangle height is proportional to the number of genes in each clade. (*B)* substitution rate (relative to mean) at each amino acid residue position for each clade, 75-residue sliding mean (gray trace). Dashed horizontal line on each plot indicates the mean evolution rate for both domains. (*C)* Bayesian posterior rate probability for LRR and RLK partitions of each clade. *** = *p*-value < .001 (unpaired Wilcoxon). (*D)* The relative evolution rate for each LRR-RLK clade, binned into 20 discrete LRR bins and 10 RLK bins, and fitted with a local polynomial regression to visualize mean rate heterogeneity.

### Positive selection is more common in LRR domains than in RLK domains

The divergent evolutionary rates we detected between LRRs and RLK domains in developmental proteins suggest divergent selective pressures acting on LRR vs. RLK domains, despite their close juxtaposition in the same protein. To determine whether LRR and RLK domains are under different selective regimes, we used our deeply sampled trees and two tests implemented in the HyPhy software suite: Mixed Effects Model of Evolution (MEME) to detect episodic diversifying (positive) selection, and Fast, Unconstrained Bayesian AppRoximation (FUBAR) to detect signatures of negative selection (Kosakovsky Pond and Frost 2005; Murrell et al. 2013). We performed these tests on our 11 LRR-RLK families as well as our outgroup *CERK1*.

MEME detected much higher rates of positive selection acting on all our LRR domains. Both MEME and FUBAR are sensitive to the number of genes included, with fewer opportunities to detect selection in clades with fewer members (Poon et al. 2009). Therefore, to normalize for different numbers of genes in each clade, we represented our results as ratios of selection pressure between each clade’s LRR and RLK domains, rather than as absolute values (fig. 2B and C). As expected (Bishop et al. 2000; X.S. Zhang et al. 2006), we found that the *CERK1* clade has a strong bias for positive selection in sequence encoding its extracellular (LysM) domain (∼41% of LysM domain residues vs ∼6% of RLK residues, ∼6.8-fold difference) (fig. S1A). Across all 11 LRR-RLK sub-clades, positive selection was more frequent in gene regions encoding LRR domains (∼10% of residues) compared to RLK domains (4% of residues) (χ^2^ test, *p*-value < 0.001) (fig. 2B). Although the absolute rate of positive selection hits varied considerably between clades, this bias towards LRRs was nonetheless true in every sub-clade (fig. 2B). For example, 11.2% of LRR domain residues and ∼4.6% of RLK residues in CLV1 were under positive selection, a ∼2.4-fold difference (fig. 2A). Interestingly, the defense proteins FLS2 and PEPR1 did not show the strong bias for positive selection acting on the LRR domain that was characteristic of CERK1. Instead, FLS2 and PEPR1 were very similar to the developmental proteins in this assay (fig. 2B, fig. S1). This suggests similar evolutionary pressures acting on developmental and defense LRR-RLKs.

**Figure 2:**
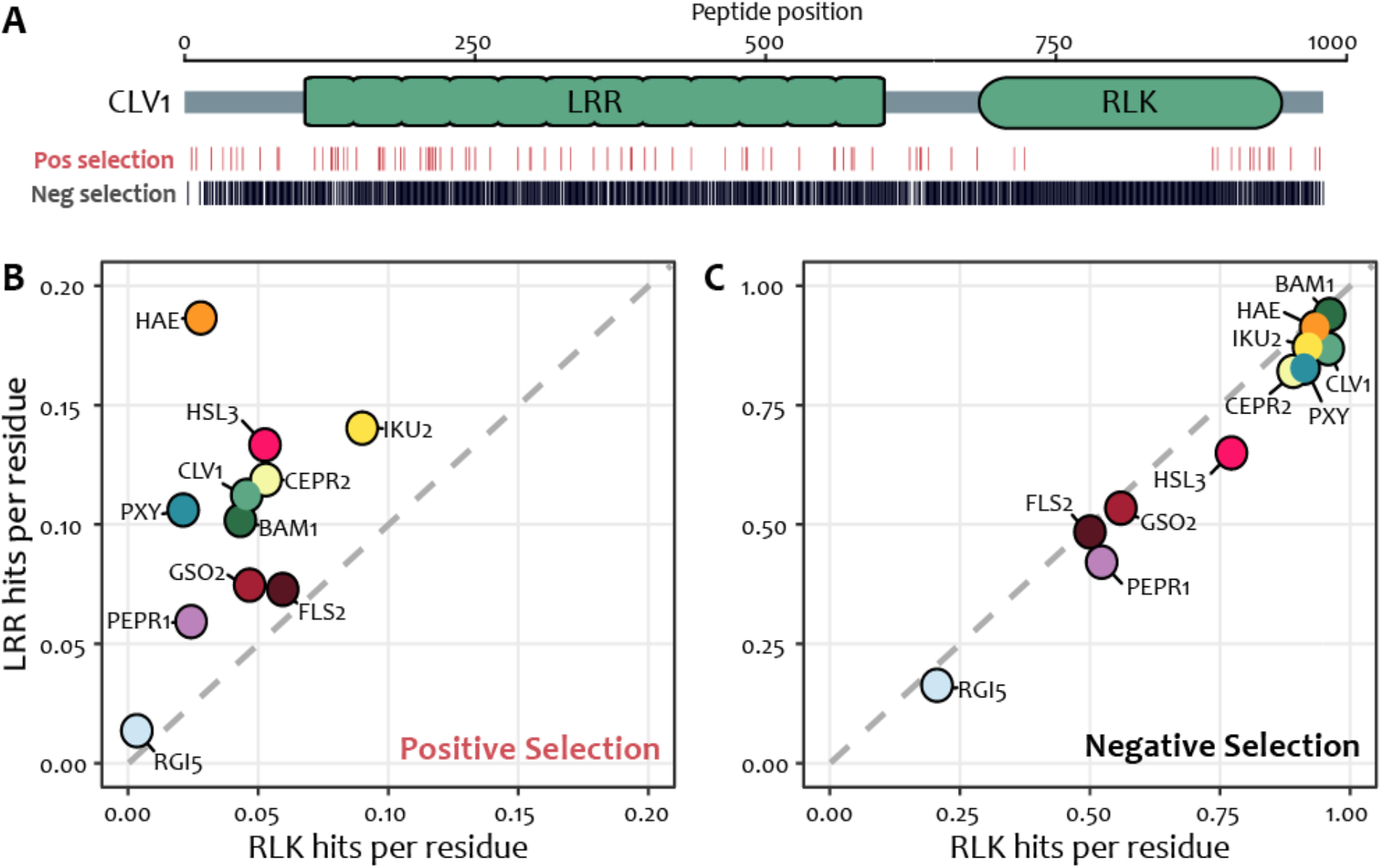
Signatures of selection are unequally distributed between LRR and RLK domains. *(A) CLV1*, as an example, with the locations of positive (red bars) and negative (black bars) selection signatures shown. *(B)* Rates per residue of signatures of positive selection in RLK (x-axis) vs LRR (y-axis) domains in each clade. *(C)* Rates per residue of signatures of negative selection in RLK (x-axis) vs LRR (y-axis) domains in each clade. Gray dashed line defines the region of no ratiometric bias between domains.

The higher positive selection acting on LRR vs. RLK domains was occurring against a background of pervasive negative selection. For example, in CLV1, nearly all residues (∼87% of LRR residues and ∼96% of RLK residues, a ∼1.1-fold difference) were under negative selection (fig. 2A). This was expected, as *clv1* mutants have severe cell proliferation phenotypes in diverse species, consistent with *CLV1* being a conserved developmental gene, and thus under strong purifying selection (Clark et al. 1993; Bommert et al. 2005; Xu et al. 2015; Whitewoods et al. 2018). This pattern extended to every clade of LRR-RLKs that we analyzed (fig. 2C). Negative selection is found in the majority of all residues for both domains (∼74% of RLK residues overall vs ∼66% of LRR residues overall), and this difference was not significant for any single clade (χ^2^ test, *p*-value > 0.05) (fig. 2C). The pervasive negative selection we detected indicates that the structures (and functions) of LRR domains on the whole are under strong evolutionary constraint, even while they undergo positive selection at certain sites. Notably, these features of LRR-RLK evolution appear to be independent of proteins’ diverse roles in development or defense. This implies that a universal mechanism is responsible for this asymmetry, which is not explained by intrinsic differences in domain leniency or defense arms-races.

### Transcriptome sequencing bias does not explain asymmetric evolutionary dynamics

One concern we had was whether qualitative differences in transcriptome sequencing could explain part or all of our results. Most genes in our analyses are from transcriptomes sequenced in the 3’ to 5’ direction (One Thousand Plant Transcriptomes Initiative 2019). LRR domains are encoded in the 5’ portions of each transcript, therefore we wondered if sequencing errors, which preferentially cluster in distal 5’ sequence, could explain the appearance of asymmetric positive selection. To check that sequencing error bias was not responsible for the elevated rate of positive selection detection in LRR domains, we performed twin analyses of the PHLOEM INTERCALATED WITH XYLEM (PXY) clade using 42 sequences each: one set translated from 1KP transcriptome sequences and a second translated from whole genome assemblies that are not directionally biased. These analyses found an identical number of positive selection hits in both domains (25 in LRRs vs. 2 in RLKs) (fig. S1). This indicates that transcriptome sequencing bias likely does not explain the consistently elevated ratio of positive selection detected in LRR domains relative to RLK domains.

### Changes in gene expression and in domain function have both likely contributed to functional divergence of LRR-RLKs

Divergent evolutionary pressures acting on LRR and RLK domains suggest that the two domains have independently diverged in function. To evaluate this prediction, we next designed *in planta* experiments to assess functional conservation of divergent LRR-RLK domains. We transformed chimeric LRR-RLKs with swapped domains into arabidopsis *clv1* single mutants and *hae; hsl2* double mutants (Scholl and Anderson 1994; Nimchuk et al. 2011). *HAE* and *HSL2* are close paralogs, and are functionally redundant with respect to floral organ abscission (Cho et al. 2008). Therefore, we will refer to these genes as *HAE/HSL2* from here on. *CLV1* and *HAE/HSL2* are distantly related LRR-RLK genes within clade XI (Man et al. 2020), separated from each other by ∼551 million years of evolution (fig. 3A, fig. S2). Over this deep time, *CLV1* and *HAE/HSL2* have diverged considerably in function. *HAE* and *HSL2* regulate cell separation at the abscission zone of sepals, petals, and stamens. *hae;hsl2* double mutants fail to shed these floral organs, which remain attached to pedicels (tan shapes at base of silique, right side of fig. 3B) (Jinn et al. 2000; Kumpf et al. 2013). *CLV1* controls proliferation of stem cells in meristems, and *clv1* mutants produce fasciated siliques with extra carpels (shorter, bulkier silique, left side of fig 3B) (Clark et al. 1993; Clark et al. 1997). Importantly, the mutant alleles we used (*clv1-15, hae-3/hsl2-3*) are recessive, and produce clear, quantifiable phenotypes (Clark et al. 1993; Jinn et al. 2000), allowing us to ask whether divergent molecular evolution was mirrored by divergent functional evolution of LRR and RLK domains.

**Figure 3:**
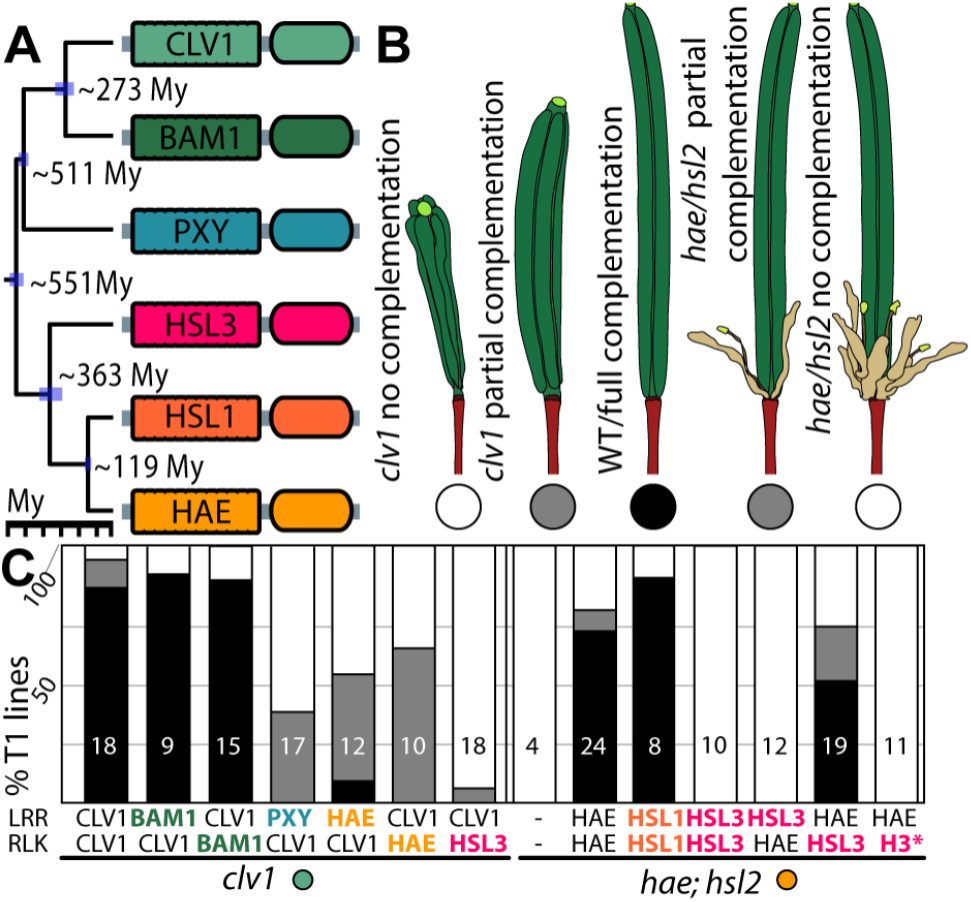
RLK and LRR domains have distinct evolutionary trajectories. Chimeric gene constructs were transformed to mutant *A. thaliana* plants to measure phenotypic rescue strength. *(A)* time-calibrated chronogram of six *A. thaliana* LRR-RLK genes showing approximate divergence times between paralogs. *(B)* Illustrations of representative arabidopsis siliques from phenotypic assays. *(C)* Complementation percentage of *clv1* or *hae/hsl2* mutants by various paralogous domains. Numbers in bars indicate independent events.

We first assessed the conservation of function between *CLV1, HAE*/*HSL2*, and their closest respective paralogs. *CLV1* is most closely related to the *BAM* genes, which are in a clade separated from *CLV1* by ∼273 million years of evolution (fig. 3A, fig. S2) (Man et al. 2020). Despite this deep divergence time, *BAM* genes also function in stem cell proliferation, and can partially compensate for the loss of *clv1* ortholog function through transcriptional regulation (Nimchuk 2017; Rodriguez-Leal et al. 2019). Consistent with this, *CLV1*^*pro*^*::BAM1*^*LRR*^*:CLV1*^*RLK*^ fully rescued the *clv1* phenotype in 8/9 events (89%), while the reciprocal swap (*CLV1*^*pro*^*::CLV1*^*LRR*^*:BAM1*^*RLK*^) fully complemented *clv1* phenotype in 13/15 events (87%), indicating strong functional conservation between the LRR and RLK domains encoded by *CLV1* and *BAM1. HAE/HSL2* and their closest paralog *HSL1* are separated by only 119 million years of evolution (fig. 3A, fig. S2), but regulate distinct processes: *HSL1* regulates stomatal development, and native *HSL1* is neither necessary nor sufficient to drive *hae; hsl2*-regulated floral organ abscission (Cho et al. 2008; Qian et al. 2018). Despite this divergence in *HSL1* function, and similar to our result for *BAM1* and *CLV1*, the full *HSL1* coding sequence driven under the *HAE* promoter can complement *hae; hsl2* double mutants. The *HAE*^*pro*^*::HSL1*^*LRR*^*:HSL1*^*RLK*^ transgene fully complemented the *hae; hsl2* phenotype in 7/8 events (88%) (fig. 3C). These data indicate that the failure of the endogenous *HSL1* locus to trigger abscission in *hae;hsl2* mutants (Cho et al. 2008; Stenvik et al. 2008) is likely due to *HSL1*’*s* divergent expression pattern (Cho et al. 2008), not the functional evolution of the HSL1 protein (Roman et al. 2022). Our results indicate that functional differences between *CLV1* and *BAM1*, and between *HAE/HSL2* and *HSL1*, are not due to changes in protein function, despite large differences in the divergence times between *CLV1, HAE/HSL2*, and their closest paralogs.

We next assessed complementation of the *hae/hsl2* double mutant with domains encoded by *HSL3*, a gene separated from *hae/hsl2* by ∼363 million years of evolution. The full *HSL3* coding sequence, expressed under the *HAE* promoter, failed to complement the *hae;hsl2* phenotype (0/10 events complemented), as did the chimeric transgene that had the LRR domain encoded by *HSL3* and the RLK domain encoded by *HAE* (*HAE*^*pro*^*::HSL3*^*LRR*^*:HAE*^*RLK*^, 0/12 events complemented) (fig. 3C). However, the reciprocal domain swap with the LRR domain encoded by *HAE* and the RLK domain encoded by *HSL3* (*HAE*^*pro*^*::HAE*^*LRR*^*:HSL3*^*RLK*^) fully complemented the *hae; hsl2* phenotype in 9/19 events (47%), and partially complemented the *hae; hsl2* phenotype in another 4/19 events (21%), suggesting substantial functional conservation of the HAE/HSL2 and HSL3 RLK domains (fig. 3C). To verify that catalytic activity of the RLK domain encoded by *HSL3* was required for complementation, we created a catalytically dead variant by introducing a single amino acid change in the ATP binding pocket (K714E) (Taylor et al. 2016). This variant (*HAE*^*pro*^*::HAE*^*LRR*^*:HSL3*^*RLK(K714E)*^) failed to complement the *hae;hsl2* phenotype (0/11 events, fig. 3C), indicating that the conserved catalytic activity of the HSL3 RLK domain is necessary and sufficient for its partial replacement of HAE/HSL2. Taken together, these data are consistent with our bioinformatic findings (fig. 1, fig. 2), and indicate that the LRR and RLK domains encoded by *HAE/HSL2* and *HSL3* have had different trajectories of functional divergence. Their LRR domains have diverged considerably and can no longer replace each other, while their RLK domains retain some conserved functionality.

Last, we assessed the ability of LRR and RLK domains encoded by deeply divergent paralogs to complement our focal mutants. We assessed complementation of *clv1* with domains encoded by *PXY* (∼511 million years of divergence), and *HAESA-LIKE3 (HSL3)* (∼551 My of divergence) (Fisher and Turner 2007). Surprisingly, RLK and LRR domains were weakly conserved in function over these very deep divergence times. The LRR domain encoded by *PXY*, together with the RLK domain encoded by *CLV1* (*CLV1*^*pro*^*::PXY*^*LRR*^*:CLV1*^*RLK*^), partially complemented the *clv1* phenotype in 6/17 (35%) transgenic lines (fig. 3C). Similarly, both the LRR and RLK domains encoded by *HAE*, together with the reciprocal CLV1 domains, partially complemented the *clv1* phenotype. Specifically, *CLV1*^*pro*^*::HAE*^*LRR*^*:CLV1*^*RLK*^ fully complemented the *clv1* phenotype in 1/12 (8%) events and partially complemented in a further 5/12 (42%) events. *CLV1*^*pro*^*::CLV1*^*LRR*^*:HAE*^*RLK*^ partially complemented in 6/10 (60%) events. Although *HSL3* is just as distantly related to *CLV1* as *HAE* is, the RLK domain encoded by *HSL3* showed very weak complementation of *clv1* when fused to the LRR domain encoded by *CLV1*. The *CLV1*^*pro*^*::CLV1*^*LRR*^*:HSL3*^*RLK*^ partially complemented the *clv1* phenotype in only 1/18 (6%) events. Together, these data suggest that some LRR and RLK domains retain vestiges of conserved function over very deep divergence times, and that conservation of domain function cannot be predicted by divergence times alone.

## Discussion

Sequences encoding the LRR domains of defense LRR-RLKs evolve under accelerated evolution with increased directional selection relative to their cognate RLK domains, presumably as a result of adapting to ever-changing pathogen effectors (Tang et al. 2010; Fischer et al. 2014; Fischer et al. 2016; Dufayard et al. 2017). Here, we show that this disjunct molecular evolution between LRR and RLK domains extends to developmental LRR-RLKs. Sequences encoding LRR domains evolve consistently faster than their RLK counterparts (fig. 1). This accelerated rate was not due to less stringent purifying selection; we detected signatures of purifying selection at about equal rates in both domains (fig. 2C), but greater rates of positive selection acting on LRR domains (fig. 2B). In contrast to the clear signal emerging from the bioinformatic analyses, our assessments of LRR and RLK domain function revealed more complexity than predicted by phylogenetic distance and evolutionary rate (fig. 3). These assays suggest that changes to both *cis-*regulation and protein structure and function were critical in the evolution of developmental LRR-RLKs.

What principle underlies the asymmetric evolution of these two domains? Some LRR-RLKs have dual roles in defense and development, and there may be undiscovered cross-talk driving some of this asymmetry (Lal et al., 2018; Verma et al., 2022; Wang et al., 2017). However, we argue that the consistency of this finding across many subclades with diverse functions suggests that asymmetric domain evolution is a widespread feature of LRR-RLKs, independent of function. Between clades of developmental LRR-RLKs, there is clearly divergent functional evolution of LRR domains: different clades of LRR-RLKs differ in function and bind different ligands, but activate intracellular signaling cascades by phosphorylating similar proteins (Zheng et al. 2019; Zhu et al. 2019; Liu et al. 2022). However, proteins within individual LRR-RLK clades often bind homologous ligands, and function in similar processes in deeply divergent lineages (Cho et al. 2008; Hirakawa et al. 2019; Rodriguez-Leal et al. 2019; Wang et al. 2019). For example, CLV1 orthologs bind CLE peptides in bryophytes, maize, arabidopsis, and tomato, lineages separated by up to ∼500 million years of evolution (Bommert et al. 2005; Hirakawa et al. 2019; Rodriguez-Leal et al. 2019). Therefore, within clades, the asymmetric evolution of LRR vs. RLK domains is more difficult to explain. It may be that LRR domains are subject to fewer structural or functional constraints than RLK domains, though to our knowledge, there isn’t evidence for this (Echave et al. 2016; Hohmann et al. 2017). Our assays also revealed pervasive negative selection occurring across both LRR and RLK domains (fig. 2C). Signatures of pervasive negative selection, higher positive selection acting on LRR domains, and variable complementarity between domains from the same gene, instead suggests asymmetry in the functional evolution of LRR and RLK domains, both between and within clades of LRR-RLK proteins.

Between clades of LRR-RLKs, escape from adaptive conflict in LRR domains may be driving asymmetric functional evolution. *Cis*-regulatory evolution can deploy conserved regulatory modules to additional, new contexts (Carroll 2008; Kramer and Li 2017). However, the dual use of a single gene in multiple contexts may incur adaptive conflict, in which genetic adaptations beneficial in one context are detrimental to another, constraining evolution (Hittinger and Carroll 2007; Flagel and Wendel 2009). Escape from adaptive conflict is the release of this constraint through gene duplications, followed by coding sequence subfunctionalization of one or both genes, allowing each gene to specialize in its respective function (Ohno 1970; Hughes 1994; Vogel and Chothia 2006). LRR-RLKs bear the hallmarks of a gene family in which escape from adaptive conflict drove functional evolution: LRR-RLKs regulate diverse developmental processes, recurrent gene duplication drove expansion of the gene family, and as we show here, they contain signatures of adaptive evolution, even in genes with developmental roles (Lehti-Shiu et al. 2012; Fischer et al. 2016; Dufayard et al. 2017; Man et al. 2020). In addition, divergent LRR-RLKs can share conserved outputs, but have divergent inputs (Zheng et al. 2019; Zhu et al. 2019; Liu et al. 2022). Prior to gene duplication, there may be low adaptive conflict in RLK domains, but high adaptive conflict in LRR domains. After gene duplication, constraint is reduced allowing accelerated optimization of formerly conflicted LRR domains, leaving the other RLK domains largely conserved in function.

Results from our domain swap experiments with HAE/HSL2 and HSL3 are consistent with functional divergence through escape from adaptive conflict in single (LRR) domains. HAE/HSL2 and HSL3 are paralogs that control unrelated developmental functions – HAE and HSL2 regulate floral organ abscission (Jinn et al. 2000), while HSL3 regulates stomatal closure (Liu et al. 2020; Liu et al. 2022). The LRR domains of HAE and HSL2 bind IDA peptides, but the LRR domain of HSL3 binds structurally distinct, non-homologous peptides called CTNIPs (Kumpf et al. 2013; Rhodes et al. 2022). In contrast to this divergent LRR binding, the RLK domains of HAE/HSL2 and HSL3 phosphorylate conserved downstream targets such as MPK3 and MPK6 (Zhu et al. 2019; Liu et al. 2022; Rhodes et al. 2022). The HSL3 LRR domain, in combination with the HAE RLK domain, failed to complement the *hae; hsl2* mutant, but the HSL3 RLK domain did complement *hae;hsl2* in the reciprocal domain swap (fig. 3C). Together, these data argue that the outputs of the HAE/HSL2 and HSL3 cytoplasmic signaling modules have not appreciably diverged in function since their duplication ∼363 Mya, but that their inputs have. Lower positive selection acting on the RLK domains of HAE/HSL2 and HSL3, conserved downstream targets, and functional conservation suggests low adaptive conflict in the output domain of the last common ancestor of these proteins. In contrast, high levels of positive selection acting on their LRR domains, nonconserved upstream triggers, and nonconservation of function indicates that diversification of the input was somehow beneficial to fitness, perhaps because of escape from adaptive conflict.

Despite a clear signal of asymmetric domain evolution emerging from our bioinformatics experiments, our heterologous transformation experiments with genes other than *HAE/HSL2* and *HSL3* revealed more complexity. When expressed under the *CLV1* or *HAE* promoters, *BAM1* and *HSL1* are each capable of performing the functions of their close paralogs, *CLV1* and *HAE*, respectively (fig. 3C). Therefore, divergence in developmental function between *BAM1* and *CLV1*, and between *HAE/HSL2* and *HSL1*, is likely not because of changes in encoded protein function, but because of divergent gene expression patterns (DeYoung et al. 2006; Cho et al. 2008; Nimchuk 2017; Qian et al. 2018; Rodriguez-Leal et al. 2019; Rhodes et al. 2022). Domains encoded by *PXY, HAE*, and *HSL3*, in chimeric transgenes with cognate *CLV1* domains, partially complement *clv1*, despite the distant relationships between *CLV1* and *HAE, HSL3*, and *PXY* (fig. 3C). Partial complementation with distantly related LRR domains could be explained by overlapping use of coreceptors, or low affinity binding of a different ligand (Hohmann et al. 2018; Qian et al. 2018; Rodriguez-Leal et al. 2019; Zheng et al. 2019; Roman et al. 2022). Some LRR domains can bind “alternate” ligands with lower affinity and serve a buffering role to regulate processes, a feature that is unlikely to be detected using their single gene mutants (Qian et al. 2018; Rodriguez-Leal et al. 2019; Roman et al. 2022). Partial complementation with RLK domains may indicate action through common intracellular signaling pathways (Meng et al. 2016; Lee et al. 2019; Zheng et al. 2019). Therefore, despite the clarity in bioinformatic signals, real-world complexities in the signaling system may obscure profound functional evolution in single gene *in planta* models.

Higher evolutionary rates and higher levels of positive selection suggest that LRR function is also changing faster than RLK function within clades of LRR-RLKs. This molecular evolution may be subtly modulating protein function. However, extensive divergence in ligand binding is unlikely to explain the divergence in LRR sequence within clades. Although ligand-receptor pairs have not been confirmed beyond arabidopsis for most LRR-RLKs, receptor-ligand relationships are deeply conserved in at least the CLV1 and HAE clades (Clark et al. 1993; Jinn et al. 2000; Bommert et al. 2005; Santiago et al. 2016; Whitewoods et al. 2018; Wang et al. 2019). In addition, key residues that mediate receptor-ligand interactions are deeply conserved within LRR-RLK clades (Santiago et al. 2016; Hohmann et al. 2018). Despite this deep conservation, receptor-ligand relationships are rarely one-to-one (Deyoung and Clark 2008; Müller et al. 2008; Ou et al. 2016; Je et al. 2018; Qian et al. 2018; Rodriguez-Leal et al. 2019). The changes to LRR domains we detected may modulate precisely which ligands bind particular LRR domains, or modulate ligand-binding kinetics. In addition, interactions with co-receptors, often mediated by LRR domains (Cui et al. 2018; Hohmann et al. 2018), can modulate receptor trafficking to and from the membrane, and the outcomes of ligand-receptor binding (Nimchuk et al. 2011; Je et al. 2018; Qi et al. 2020). Thus, while the function of particular LRR-RLKs may be broadly conserved over deep time, rapid evolutionary change and positive selection may indicate incremental and subtle change to protein (and LRR domain) function.

Our work has implications for agriculture and crop engineering, where the subtle modulation of plant traits was critical in domestication, and is becoming useful in crop improvement using genome engineering. This subtle modulation is often accomplished through *cis-*regulatory change (Meyer and Purugganan 2013; Rodríguez-Leal et al. 2017; Stitzer and Ross-Ibarra 2018; Bartlett et al. 2022). However, coding changes have been important in plant evolution and development (Bartlett and Whipple 2013; Bartlett 2019; Bartlett et al. 2022). Coding sequence change could be more easily implemented and may have more predictable outcomes than *cis*-regulatory sequences, which are incompletely understood in plants (Rodríguez-Leal et al. 2017; Marand et al. 2023). Importantly, the LRR-RLK protein family regulates many aspects of plant development that are important agronomically, and have already been targeted in crop improvement efforts (Bommert et al. 2005; Xu et al. 2015; Je et al. 2016; Yang et al. 2018; Shao et al. 2019). Deep knowledge of LRR-RLK molecular biology in arabidopsis, and its powerful molecular toolbox, allows for the detailed dissection of how residues under selection may be impacting protein function over deep time (Bartlett 2019). Indeed, selection tests pinpointed residues that mediate protein-protein interactions between B-class transcription factors, interactions which affect protein degradation and floral organ development (Bartlett et al. 2016; Abraham-Juarez et al. 2020). Our assessments of selection acting on key LRR-RLK residues, coupled to deep knowledge of LRR-RLK biology, could provide novel solutions in crop improvement.

## Materials and Methods

### Sequence retrieval and curation

For each clade, we collected gene locus identifiers (Man et al. 2020), and peptide sequences obtained from the primary transcript annotation databases for *Arabidopsis thaliana, Amborella trichopoda, Brachypodium distachyon, Oryza sativa* (rice), *Solanum lycopersicum* (tomato), *Populus trichocarpa* (poplar), *Selaginella moellendorffii*, and *Physcomitrella patens*, and *Zea mays* (maize) from Phytozome ver. 12 (Goodstein et al. 2012). We aligned these sequences using *MAFFT* v. 7.313 (Katoh and Standley 2013) and built a Hidden Markov Model (HMM) profile using *HMMER* v. 3.1b2 (Altschul et al. 1990; Eddy 2011). We used this HMM profile to scan every peptide database from the One Thousand Plants Transcriptome Initiative (1KP), and the top two scoring hits from each genome were collected (One Thousand Plant Transcriptomes Initiative 2019). We then scanned hits for protein domains using *HMMER* v3.1b2 and the Pfam protein profile HMM database v31.0 using the “trusted cutoff” bit score gathering threshold (Eddy 2011; El-Gebali et al. 2019). `Only hits with detected LRR and RLK domains, and with a length ≥85% of the Arabidopsis anchor gene were included. We` aligned hits using the L-INS-i iterative refinement algorithm implemented in *MAFFT* v. 7.313 (Katoh and Standley 2013), filtered for homoplastic positions by *Noisy* v1.5.12 (Dress et al. 2008), and inferred a gene tree using *IQtree* v1.6.3 (Nguyen et al. 2015). We interpreted and visualized the tree using package *ggtree* v1.10.0 in *R* v3.4.3 (R Core Team 2017; Yu et al. 2017), and the 1KT genes falling into each target clade were iteratively used as seed sequences in another round of BLAST and HMM profiling until stable protein trees for each clade were inferred (Table 1).

### Maximum likelihood gene tree inference

For each clade, we build alignments using search hits and *AtERECTA* (AT2G26330) as the outgroup using the L-INS-i iterative refinement algorithm implemented in *MAFFT* v. 7.313 (Katoh and Standley 2013). Next we filtered for homoplastic positions using *Noisy* v1.5.12 (Dress et al. 2008), and used the alignment to find the best-fitting model of protein evolution and infer a gene tree using maximum likelihood analysis with 1000 bootstrap replicates using *IQtree* v1.6.3 (Nguyen et al. 2015). We interpreted and visualized the tree using package *ggtree* v1.10.0 in *R* v3.4.3 (R Core Team 2017; Yu et al. 2017)

### Rate of evolution by site

Because the 1KP dataset was enriched for angiosperms, and we wanted densely sampled protein trees, we trimmed each alignment and tree to include only angiosperm peptide sequences, with *Amborella trichopoda* genes positioned as sister to all other sequences in each analysis (Mathews and Donoghue 1999; Soltis et al. 2008). For each clade, we aligned peptide sequences from each clade using the L-INS-i iterative refinement algorithm implemented in *MAFFT* v. 7.313 (Katoh and Standley 2013), and used this to infer the posterior mean site evolution rate with a Yule speciation prior and a random starting tree using the -wsr function in *IQtree* v1.6.3 (Nguyen et al. 2015). To prevent alignment artifacts from distorting the mean rate calculations, we removed positions from the alignment not present in the Arabidopsis gene from which the clade is named (anchor gene). We calculated and visualized the sliding window mean-rate trace for each residue of the Arabidopsis anchor gene using a custom R script utilizing the packages Biostrings v.2.54.0 (Pagès et al. 2017), *ggplot2* v3.3.0 (Wickham et al. 2019), and zoo v1.8-7 (Zeileis and Grothendieck 2005). *LRR* and *RLK* domain coordinates for the Arabidopsis anchor gene were taken from Uniprot definitions (UniProt Consortium 2015). To calculate mean evolution rates across all clades, we binned rates for each clade into 3 N-terminal, 20 LRR, 3 transmembrane, 10 RLK, and 2 C-terminal bins, shown fitted with a local polynomial regression with an α=0.1.

### Rate of evolution by domain

For each clade alignment, LRR and RLK partitions were defined based on the Uniprot domain annotations of the clade’s Arabidopsis anchor gene (UniProt Consortium 2015). We performed Bayesian analyses using *BEAST* v2.6.1 (Bouckaert et al. 2019) on the CIPRES Science Gateway (Miller et al. 2010). In these analyses, the peptide sequence alignment is partitioned into LRR and RLK domains, and the tree and partition substitution rates are co-estimated, then weighted over tree-space posterior probabilities to give a rate distribution estimate per domain. To infer relative domain rates, we used the following parameters: Tree and clock models, but not site models, were linked during the analysis, to provide independent site rate estimates for the partitions. A gamma site model with six discrete estimated rate categories was set for each partition to ensure appropriate gamma rate heterogeneity, using a JTT amino acid substitution matrix which was found to best fit LRR-RLKs (Susko et al. 2002; Man et al. 2020). Strict and relaxed molecular clock models were compared using a nested sampling approach to generate marginal likelihoods for a Bayes Factor estimation and a likelihood ratio test (Kass and Raftery 1995; Brown and Yang 2011; Russel et al. 2019). Based on these tests we proceeded using a relaxed clock model, which was subsequently used for all further analyses. Clade trees were then inferred under a Calibrated Yule model and a Birth-death model, and had statistically identical Maximum Clade Credibility trees as calculated by *Tree Annotator* v2.6.2 (Bouckaert et al. 2019), therefore Birth-death models were used in all clades as in (Magallón et al. 2015). The node containing all major angiosperm clades other than *A. trichopoda* was calibrated to 135-137 Ma (lognormal prior distribution, 95% posterior density credibility interval) (Magallón et al. 2015). Markov chain Monte Carlo (MCMC) chains were run for sufficient length to ensure complete mixing and convergence of molecular clock rate estimate and other parameters used, thresholded by Effective Sample Sizes (ESS) > 100, measured using *Tracer* v1.6 (Rambaut et al. 2018). Within each clade, rate estimate distributions did not have equivalent variances (F-test, p < .001 for all clades) and were therefore compared using non-parametric unpaired Wilcoxon tests.

### Selection tests

We performed tests looking for signatures of selection using the nucleotide sequences corresponding to the peptide MSAs used to infer evolutionary rates. We aligned nucleotide sequences by their codon translations using the BLOSUM45 scoring matrix in *MAFFT* v. 7.313 (Katoh and Standley 2013) as implemented in *Geneious* v10.0.8 (Kearse et al. 2012). These MSAs were used to run tests in the *HyPhy* suite (Pond et al. 2005): *FUBAR* (Murrell et al. 2013), and *MEME* (Murrell et al. 2012), as implemented on the Datamonkey web server (Weaver et al. 2018). MEME hits were thresholded at the default p-value of 0 .1 and FUBAR hits were thresholded by a more stringent threshold of 0.999 posterior probability. Site-by-site selection evidence from these tests were visualized using a custom *R* script utilizing *ggplot2* v3.3.0 (R Core Team 2017; Wickham et al. 2019). The alternate *GSO2* selection tests utilized only CDS sequences from genomic assemblies available from homologs detected from Ensembl Plants (Kersey et al. 2018).

### Maximum Likelihood inference of gene tree backbone

We aligned arabidopsis, rice, tomato, and *A. trichopoda* peptide sequences from each LRR-RLK clade using *CERK* LysM-RLK as the outgroup. We used the E-INS-i algorithm in *MAFFT* v. 7.313 (Katoh and Standley 2013) due to the unrelated LysM domain of *CERK1* paralogs in the same position as LRRs. Next we filtered the alignment for homoplastic positions using *Noisy* v1.5.12 (Dress et al. 2008), and found best-fit model of protein evolution and inferred a gene tree by maximum likelihood analysis with 1000 bootstrap replicates using *IQtree* v1.6.3 (Nguyen et al. 2015). This tree was visualized using *FigTree* v1.4.3 (Bouckaert et al. 2019).

### Chronogram for inference of gene clade divergence times and tree topology

Four genes from each clade were extracted: the *A. trichopoda* gene, and a representative from rosids, asterids, and monocots, generally *A. thaliana*, tomato, and rice, and well-supported *S. moellendorffii* and *P. patens* genes that are sister to these clades. We aligned these using the L-INS-i iterative refinement algorithm implemented in *MAFFT* v. 7.313 (Katoh and Standley 2013). We used this alignment to perform Bayesian analyses using *BEAST* v2.6.1 (Bouckaert et al. 2019) on the CIPRES Science Gateway (Miller et al. 2010). This analysis was run as a single partition, using the JTT substitution matrix, and a relaxed clock model. Priors included a Yule tree model, and Most Recent Common Ancestor (MRCA) nodes: eudicot MRCA was uniform 111-131 My, angiosperm MRCA was uniform 173-199 My, tracheophyte MRCA was uniform 410-468 My, and embryophyte MRCA was uniform 465-533 My (Magallón et al. 2015; Kumar et al. 2017). MCMC chains were run for sufficient length to ensure complete mixing and convergence of molecular clock rate estimate and other parameters used, thresholded by Effective Sample Sizes (ESS) > 100, measured using *Tracer* v1.6 (Rambaut et al. 2018). The final tree was visualized using *FigTree* v1.4.3 (Bouckaert et al. 2019) (fig. S2).

### Construction of chimeric transgenes

We utilized the MoClo system of Golden-Gate molecular assembly to make full-length domain swaps constructs using sequences encoding promoters, LRR domains, RLK domains, and terminators (Engler et al. 2014). For *CLV1* promoter constructs, we replicated the promoter/5’UTR and terminator/3’UTR coordinates published previously (Nimchuk et al. 2011). For *HAE* promoter constructs, we determined the promoter/5’UTR by conservation with other species using Vista plots in Phytozome’s JBrowse (Goodstein et al. 2012) to be 1,609bp upstream the gene’s start codon, and utilized the terminator from the *nos* gene in *A. tumefaciens* (Engler et al. 2014). The sequence encoding protein domains was amplified using PCR with Col-0 gDNA template, or synthesized commercially. We assembled all transgenic constructs into either the pICH86966 T-DNA plant transformation vector from the MoClo kit (Engler et al. 2014) or a plasmid derived from pH7m24GW (O’Connor et al. 2017) and verified by Sanger sequencing or whole plasmid sequencing.

To standardize domain coordinates across genes used for domain swaps, we made a translation alignment between all genes and engineered a synthetic 4-bp Golden-Gate assembly joining overlap within the transmembrane domain that introduced only synonymous coding alterations in all of the genes used in our assay.

### Plant growth conditions and plant transformation

Seeds were sown on the surface of soilless medium in 3” pots, watered, and stratified covered in dark at 4°C for 3 days, then transferred to a growing rig (PAR ∼ 100 μmol/m^2^/s), with a 16 h light and 8 h dark cycle, constant temperature ∼20°C.

*clv1-15* (WiscDsLox489-492B1) was kindly provided by Zachary Nimchuck (Nimchuk et al. 2011). *hsl2-3/hae-3* (CS69822) seed was obtained from the Arabidopsis Biological Resource Center (Scholl and Anderson 1994). Both mutants are in the Columbia background. Into these plant lines we transfected LRR-RLK constructs using *Agrobacterium tumefaciens* strain GV3101 with the floral dip method (X. Zhang et al. 2006). Transgenic T1 seeds were selected according to (Davis et al. 2009) and transplanted from selection plates to soilless medium in pots at two weeks of age and grown to maturity.

### Phenotype scoring

All T1 plants were phenotyped as soon as the oldest 10 flowers on the primary inflorescence could be scored for abscission of floral organs (for *hae;hsl2* rescue constructs) or carpel number (for *clv1* rescue constructs). Events from *HAE* constructs were scored by lightly brushing the inflorescence twice, with a 90° turn between, through a pair of soft bristle paint brushes affixed with bristles oriented towards each other. After brushing, we assessed *hae; hsl2* double mutant complementation by binning each transformed event into one of three categories: fully complemented (all floral organs abscised), partially complemented (some abscised, some remain attached to pedicel), or not complemented (no floral organs abscised). The complementation strength for a construct as a whole was scored as a “complemented event” if <9 (out of 10) flowers were WT (partially complemented flowers count as half), or a “partially complemented event” if >1 and ≤9 flowers were wild type. Events from *CLV1* constructs were scored by counting carpels on each silique from each mature transformant (9 - 24 independent T1 lines) using a dissecting microscope, and calculating the mean carpel number per silique in each event. Events were binned into three categories: fully complemented (≤2.5 carpels per silique), partially complemented (2.5 to 3.25 carpels per silique), or not complemented (>3.25 carpels per silique). The complementation strength for a construct as a whole was calculated as the fraction of flowers across all events that were fully complemented, with partial complementation events contributing at half rate.

## Data Availability Statement

All alignments, trees, rate test results, selection test results, and phenotyping data are available at dryad (doi:10.5061/dryad.jm63xsjg5). Code for analyses is available on the Bartlett lab’s GitHub (https://github.com/BartlettLab/Domain_evolution)

## Acknowledgements

We would like to thank Courtney Babbitt, Remco Bouckaert, Heather Bracken-Grissom, Joseph Gallagher, Jason Kamilar, Zachary Nimchuk, and Devin O’Connor, for technical support, helpful discussion, reagents, and seed. Amber De Neve, Joseph Gallagher, and Erin Patterson provided helpful comments on an earlier version of the manuscript. This work was supported by the National Science Foundation (IOS-2129189 to M.B.), a Lotta Crabtree fellowship (J.M.), and a UMass Biotechnology Training Program fellowship (T. H.).

## Supplemental Figures

**Figure S1.**
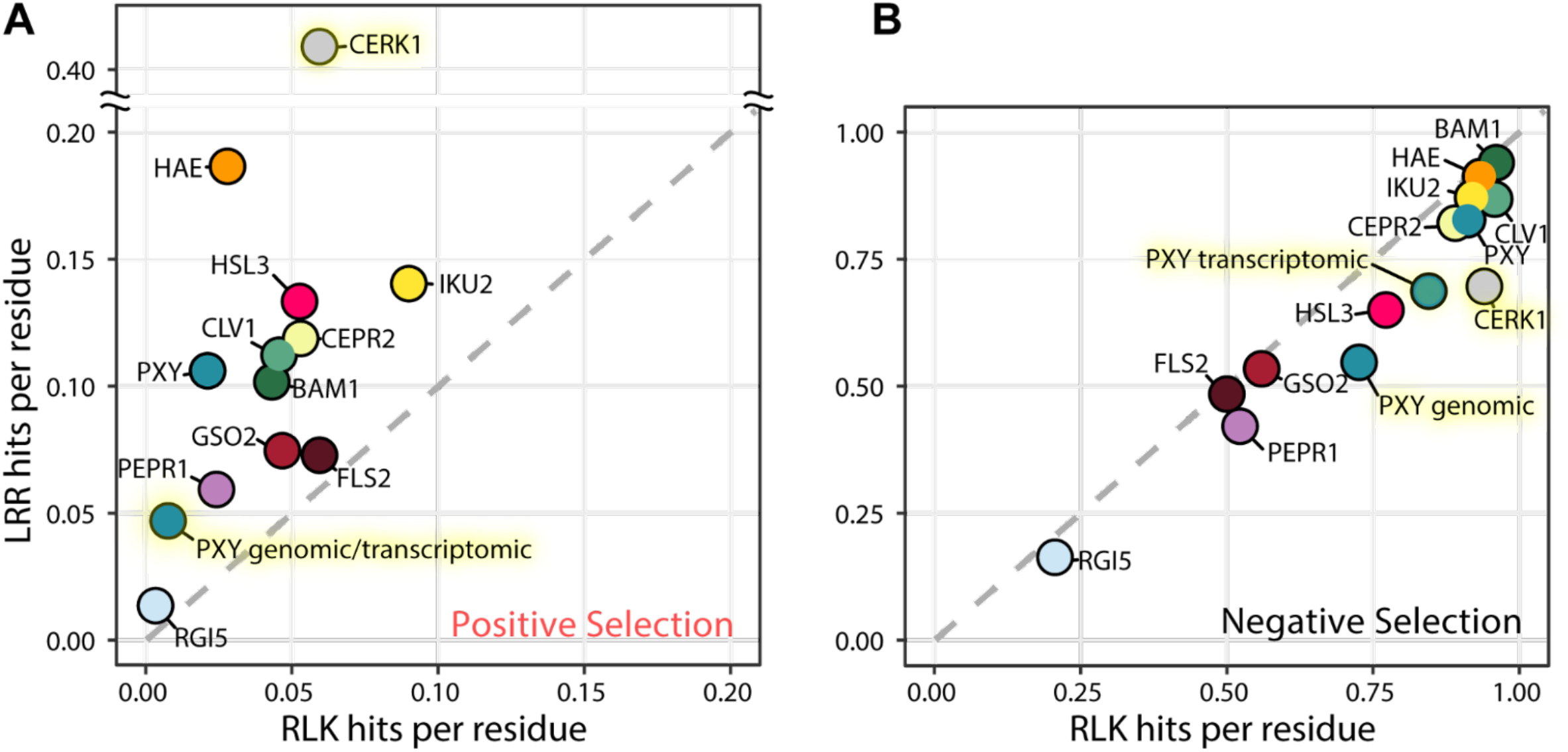
*CERK1* has an unequal distribution of selection signatures. *(A)* Positive selection detection rate for *CERK1, PXY* from genomic and *PXY* from transcriptomic datasets are also shown in addition to the other LRR-RLK clades. The two new *PXY* data points are identical. *(B)* Negative selection detection rate for *CERK1, PXY* from genomic and *PXY* from transcriptomic datasets are also shown in addition to the other LRR-RLK clades. Gray dashed line defines the region of no ratiometric bias between domains.

**Figure S2.**
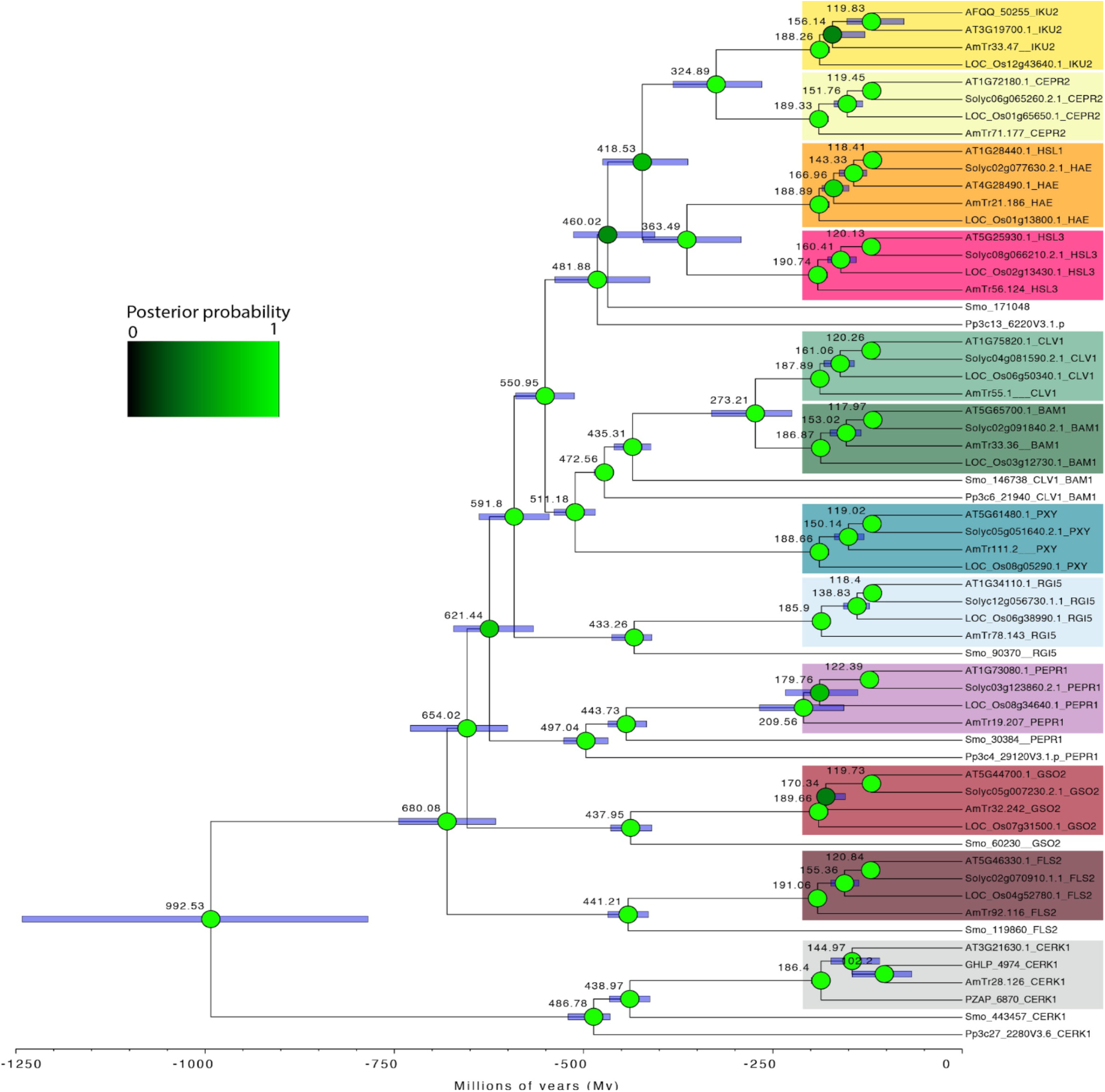
Chronogram for inference of gene clade divergence times and tree topology. 4 angiosperm representatives from 11 LRR-RLK clades, and from *CERK1*, as well as well-supported representatives from *S. moellendorffii* and *P. patens* sister to those clades, were used to infer a time-calibrated gene divergence tree by Bayesian analysis. Blue bars over nodes represent 95% distribution probability range, while circles over nodes are colored according to posterior probability. Number next to each node is the estimated mean divergence time.

